# *fluxweb*: a R package to easily estimate energy fluxes in food webs

**DOI:** 10.1101/229450

**Authors:** Benoit Gauzens, Andrew Barnes, Darren Giling, Jes Hines, Malte Jochum, Jonathan S. Lefcheck, Benjamin Rosenbaum, Shaopeng Wang, Ulrich Brose

**Affiliations:** EcoNetLab, German Centre for Integrative Biodiversity Research (iDiv) Halle-Jena-Leipzig, Deutscher Platz 5e, 04103 Leipzig, Germany; Institute of Biodiversity, University of Jena, Dornburger Str. 159, 07743 Jena, Germany; School of Science, University of Waikato, Private Bag 3105, Hamilton 3204, New Zealand; Institute of Landscape Ecology, University of Münster, Heisenbergstrasse 2, 48149 Münster, Germany; Institute of Biology, Leipzig University, Johannisallee 21, 04103 Leipzig, Germany; Institute of Plant Sciences, University of Bern, Altenbergrain 21, 3013 Bern, Switzerland; Tennenbaum Marine Observatories Network, MarineGEO, Smithsonian Institution, Edgewater, Maryland; Institute of Ecology, College of Urban and Environmental Science, Peking University, Beijing 100871, China

## Abstract

- Understanding how changes in biodiversity will impact the stability and functioning of ecosystems is a central challenge in ecology. Food-web approaches have been advocated to link community composition with ecosystem functioning by describing the fluxes of energy among species or trophic groups. However, estimating such fluxes remains problematic because current methods become unmanageable as network complexity increases.
- We developed a generalisation of previous indirect estimation methods assuming a steady state system [1, 2, 3]: the model estimates energy fluxes in a top-down manner assuming system equilibrium; each node’s losses (consumption and physiological) balances its consumptive gains. Jointly, we provide theoretical and practical guidelines to use the *fluxweb* R package (available on CRAN at https://bit.ly/2OC0uKF). We also present how the framework can merge with the allometric theory of ecology [4] to calculate fluxes based on easily obtainable organism-level data (i.e. body masses and species groups -eg, plants animals), opening its use to food webs of all complexities. Physiological losses (metabolic losses or losses due to death other than from predation within the food web) may be directly measured or estimated using allometric relationships based on the metabolic theory of ecology, and losses and gains due to predation are a function of ecological efficiencies that describe the proportion of energy that is used for biomass production.
- The primary output is a matrix of fluxes among the nodes of the food web. These fluxes can be used to describe the role of a species, a function of interest (e.g. predation; total fluxes to predators), multiple functions, or total energy flux (system throughflow or multitrophic functioning). Additionally, the package includes functions to calculate network stability based on the Jacobian matrix, providing insight into how resilient the network is to small perturbations at steady state.
- Overall, *fluxweb* provides a flexible set of functions that greatly increase the feasibility of implementing food-web energetic approaches to more complex systems. As such, the package facilitates novel opportunities for mechanistically linking quantitative food webs and ecosystem functioning in real and dynamic natural landscapes.

## 1 Introduction

In recent years, there have been multiple calls for the reconciliation of food web structure and ecosystem functioning, to better understand how changes to ecological networks will influence the stability and functioning of ecosystems [5, 6, 7]. Energetic food-web approaches can be used to quantify a key aspect of ecosystem functioning, energy flux, as a way of characterizing ecological processes that are driven by trophic interactions among nodes in food webs [8, 1,9]. As such, energy fluxes can be used to quantify functions such as herbivory or productivity. They can also be integrated into the classical framework of Lotka-Volterra equations to estimate stability [10, 11].

Despite interest in using quantitative networks [12, 13, 14], they are still rarely employed for describing natural communities. This is, in part, because quantifying interaction strengths or fluxes in food webs remains a deceptively difficult problem, often requiring intensive experimental and observational efforts. A viable solution is to use mathematical proxies for system, and/or organismal level parameters for calculating energy fluxes through networks based on easily accessible parameters, rather than attempting to measure flux through the whole network. At the system level, for example, inverse matrix reconstruction (commonly referred to as ‘ecological network analysis’) [15, 16], or the ‘food-web energetics approach’ [1, 2, 3] have gained some support. These approaches, which are both based on the same steady state assumption (i.e., populations are at equilibrium densities), require reasonable knowledge of the focal system such as network topology. However, a major difference relates to the solution provided by these two methods. The ecological network analysis produces an infinite number of solutions and requires an a posteriori selection function. In contrast, the web energetic approach assumes that fluxes are driven by a top-down effect (energetic demand of predators drive their ingoing fluxes) to guarantee a unique solution for each dataset. Previously, however, scientists using the ‘food web energetics’ approach [2, 9, 17], have manually calculated fluxes, which can become exceedingly unmanageable as the complexity of the food web increases. Therefore, there is urgent need for a generalized automation of this method. Interaction strengths can also be quantified by focusing on organism-level parameters related to the metabolic theory of ecology [4]. Generalized al-lometric approaches utilize general patterns of functional responses that depend upon body size ratios between consumers and their resources [18, 19], opening ways for determining interaction strengths in response to commonly available data such as the abundances and body masses. Allometric rules have been successfully applied to predict fluxes in simplified systems with a few species [20]. However, these results have not yet been generalized for use in complex networks.

Here we present the methodological and mathematical framework that underlies the food web energetics approach and provide theoretical and practical guidelines for using the fluxweb R package. We then show how the framework presented here can easily merge with the allometric theory to estimate energy fluxes in complex natural food webs. In doing so, we support proposals to create a framework allowing for the estimation of energy fluxes in trophic networks using widely available ecological information such as biomass, metabolic demand, ecological efficiencies or network topology [7].

## 2 The underlying model

The model underlying the food web energetics approach assumes a steady state. It implies that *L*_*i*_, the total amount of energy lost by a species *i*, either by consumption or physiological processes, is exactly compensated by the metabolized energy it gains from consumption *G*_*i*_. It will thus solve the equation

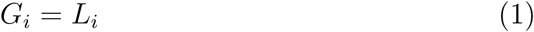

knowing that

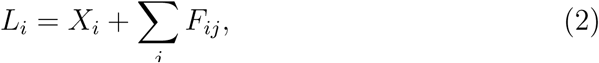

where *X*_*i*_ defines energetic losses from species *i* such as through metabolism, and *F*_*ij*_ is the flux from species *i* to its consumer species *j*. Then, gains are the part of ingoing fluxes once losses due to feeding efficiency are removed.

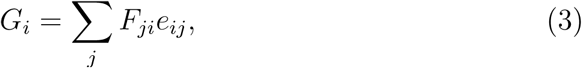

where *e* defines a species’ feeding efficiency. This parameter can either depend only on *i* (efficiencies depends only on the predator identity), only on *j* (efficiencies related to prey identity), or on both. More details about parameters can be found in section 3. Any flux *F*_*ij*_ can be written as *F*_*ij*_ = *W*_*ij*_*F*_*j*_, where *F*_*j*_ is the sum of all ingoing fluxes to species *j*, and *W*_*ij*_ defines the proportion of *F*_*j*_ that is obtained from species *i* (Σ_*i*_ *W*_*ij*_ = 1). The package offers the possibility to scale predator preferences to the distribution of prey body masses. We thus obtain the following model for determining each species’ sum of ingoing fluxes:

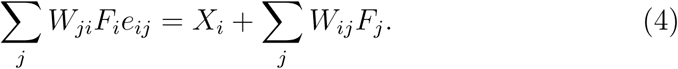

This equation is solved in two stages: first, the sum of ingoing fluxes for each species is computed. Then, individual fluxes for each pairwise predator-prey interaction are calculated using predator preferences (set in *W*).

The solution for eq. 4 depends on the chosen definition of feeding efficiency (assigned based on the predator, prey, or link identity) (see Supporting Information I for demonstrations) and is as follows:

- Efficiencies depending on predator identity

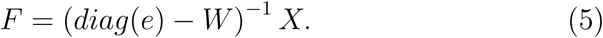 *F* is the vector such that *F*_*i*_ describes the sum of ingoing fluxes to species *i*, *e* is the vector of feeding efficiencies, such that *e*_*i*_ describes the efficiency of a predator *i* (see related paragraph in section 3 for more precise definitions of feeding efficiencies) with *e*_*i*_ = 0 if *i* is basal. *W* is the matrix such that *W*_*ij*_ sets the proportion of ingoing fluxes to species *j* from species *i* and *X* is the vector defining the sum of energetic losses for each species.
- Efficiencies depending on prey identity

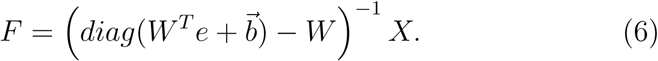 In this case, *e* is the vector such that *e*_*i*_ expresses a prey-related efficiency. 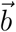 is a vector such that *b*_*i*_ is equal to 1 if species *i* is basal, 0 if it is not basal. The addition of this last vector is needed to solve the system. Ecologically, it simulates the addition of a nutrient node on which all basal species feed with an efficiency of 1.
- Efficiencies depending on link identity (both prey and predator)

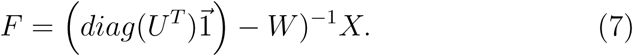 Here, 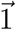 is a vector of ones, *U* is the matrix defined by the element-wise multiplication of matrices *W* and *e*: *U*_*ij*_ = *W*_*ij*_*e*_*ij*_. *e* is a matrix such that *e*_*ij*_ is the assimilation efficiency of species *j* feeding on species *i*.

## 3 Defining the parameters

A great advantage of the food web energetics method is that it offers a flexible quantitative framework that can be used to test many different ecological hypotheses related to fluxes in networks [21, 3]. Parameters used to configure the model can be taken from the literature, estimated from direct field measurement or assessed from general scaling relationships using easily accessible species (e.g., body size) and/or environmental (e.g., temperature) information. Therefore, the *fluxweb* package is a tool that is highly applicable for both experimental/empirical approaches aiming to describe natural systems and for theoretical approaches requiring generic solutions.

In the following section, we will describe the different parameters needed and how they can be estimated (see table 1 for examples).

**Table 1:**
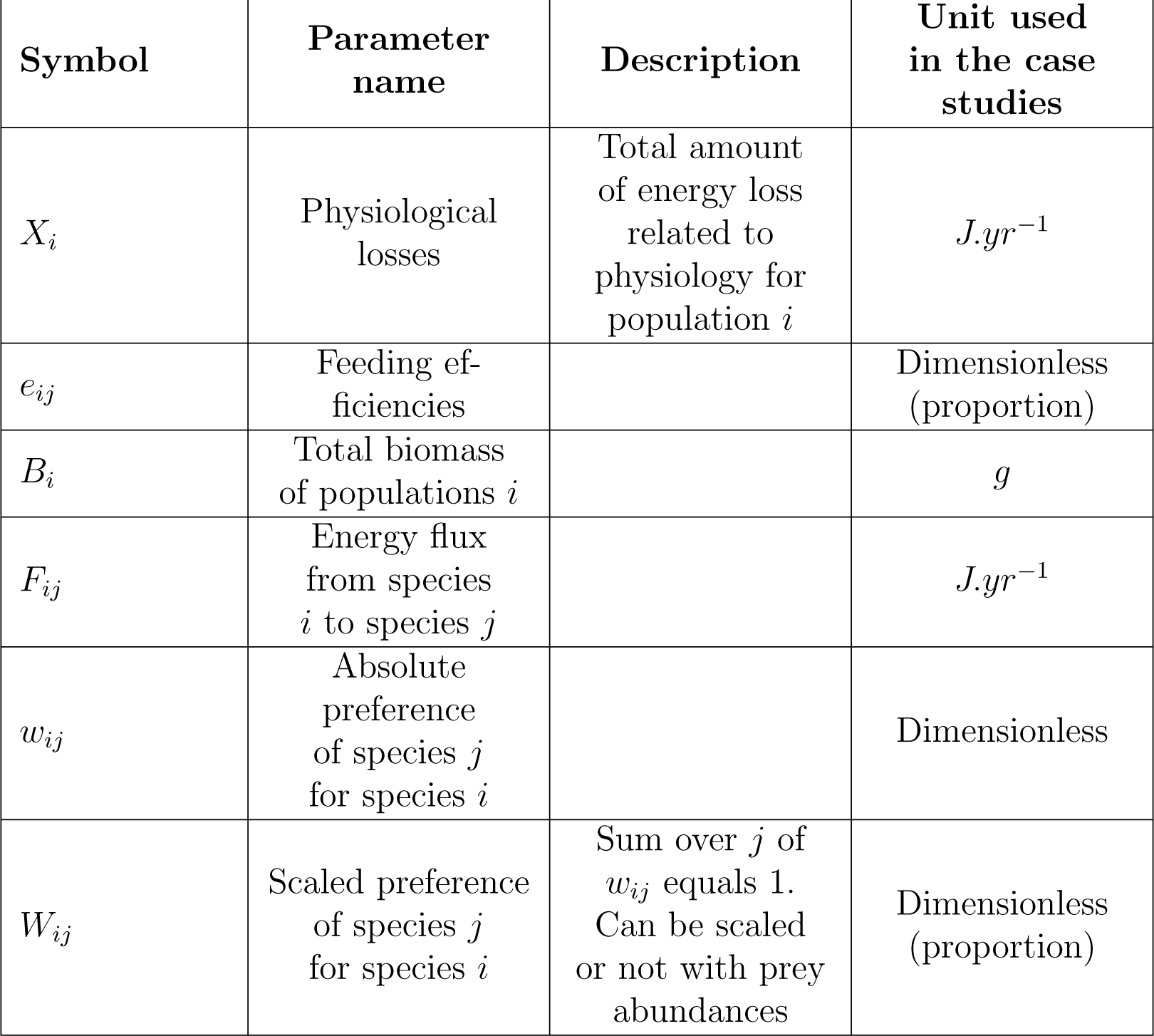
Description of the different parameters used in the *species.level* and *group.level* case studies and their meaning. The units are only examples and might depend on user choices, as long as global coherence is preserved.

| Symbol | Parameter name | Description | Unit used in the case studies |
| --- | --- | --- | --- |
| $X_i$ | Physiological losses | Total amount of energy loss related to physiology for population $i$ | $J.yr^{-1}$ |
| $e_{ij}$ | Feeding efficiencies | | Dimensionless (proportion) |
| $B_i$ | Total biomass of populations $i$ | | $g$ |
| $F_{ij}$ | Energy flux from species $i$ to species $j$ | | $J.yr^{-1}$ |
| $w_{ij}$ | Absolute preference of species $j$ for species $i$ | | Dimensionless |
| $W_{ij}$ | Scaled preference of species $j$ for species $i$ | Sum over $j$ of $w_{ij}$ equals 1. Can be scaled or not with prey abundances | Dimensionless (proportion) |

### Physiological losses (*X*_*i*_)

Depending on user assumptions and choices, different ecological processes can be used. Classical choices are often:

- Metabolic rates [4]
- Death rates [22]
- Potentially more complex allometric functions, including time allocated to resting or hunting and associated energy costs [23]

Metabolic rates and death rates can be measured for simplified systems such as microcosms experiments [24]. If the complexity of the network considered prevents such measurements, they can be estimated for each taxonomic group *i* using the classic allometric equation [4]

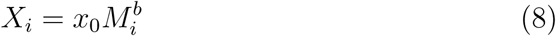

where *X*_*i*_ represents a parameter related to the physiology of species *i*. *x*_0_ and *b* are constants associated with parameter *X*_*i*_ and *M*_*i*_ is body mass. References for the choice of constant values associated with each model parameter can be found in the descriptions below. Depending on the amount of ecological information available, or precision required, parameters *x*_0_ and *b* can be quite general (i.e., the same value for all species), or more specific (i.e., applying one parameter value per functional group, taxonomic group, or species [25]). As *X*_*i*_ is typically estimated per unit biomass, setting the correct value for estimating energy flux is done by a simple multiplication by a species’ total biomass. It is interesting to note that the loss parameter can be used to drop the steady state assumption. Indeed, with two or more samples of the same system at different dates, it is possible to add the biomass differences observed as positive (i.e. loss of biomass on time) or negative (i.e. gain of biomass in time) energetic losses after a conversion in coherent units. Removing the equilibrium assumption however prevents the use of the stability functions (as they are defined only for steady state systems).

### Efficiencies (*e*)

*fluxweb* offers the possibility to use a variety of input parameters that define energetic losses, for which different aspects of ecological efficiency must be employed. If metabolic rate is used to parametrise energetic loss, then assimilation efficiency must also be provided (i.e., the proportion of consumed energy that is assimilated for respiration and biomass production). If mass-specific death rates are used in place of metabolism (sensu [21]), users should use the product of assimilation efficiency and production efficiency (percentage of assimilated energy that is used for biomass production). The *fluxweb* package offers three different options for defining ecological efficiencies: consumer-defined, resource-defined, or link-defined (considering both predator and prey identity) efficiencies. These options correspond respectively to the values *pred, prey* and *link.specific* for the *ef.level* argument. If, within a single study, each consumer has a relatively homogeneous resource pool (i.e., consumers are trophic specialists such as strict herbivores or strict carnivores), defining efficiencies at the consumer level could be the standard option. However, if a single consumer node draws on a variety of resource nodes (e.g., plants, detritus and animals), efficiencies can be defined at the resource level to account for differences in resource quality ingested by a consumer species. For this last approach, efficiency values that relate to the different groups of organisms can be found in the literature [26].

### Preferences (*W*)

Preferences depict the feeding behavior of predator species and should quantify their foraging choices. Depending on system and user choice, they can be absolute preferences or per capita. The package offers the possibility to estimate or scale preferences using a linear scaling with prey biomass:

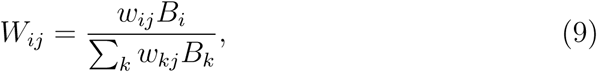

where *B*_*i*_ sets the biomass of species *i* and *w* is defined by a priori preferences from users. *w* values are values from the network adjacency matrix (i.e. the matrix such that the value of the *i*^*th*^ line and the *j*^*th*^ column is non zero if predator *j* feeds on prey *i*). Thus, preferences can be defined as a combination of active choice only (by setting the *bioms.pref* option to *FALSE* and providing preferences as values in the network adjacency matrix), relative availability of prey only (by setting the *bioms.pref* option to *TRUE* and providing a binary adjacency matrix for the network) or a combination of both, if preferences values are provided in the adjacency matrix and the option *bioms.pref* is set to *TRUE*.

### Species biomasses (*B*)

Biomasses are used (depending on user choices) to scale losses (if they are provided per biomass unit) and preferences. It is therefore an optional parameter.

## 4 *fluxweb* functionalities

Above we presented the theoretical background used by *fluxweb* to determine fluxes in food webs with the *fluxing* function. However, the package offers several other possibilities. Under the steady state assumption, it is quite straightforward to relate estimated fluxes to the equilibrium state of a set of ordinary differential equations depicting population dynamics (Lotka-Volterra systems of equations). This offers the possibility to gain insight into network stability using the methods established for such equation systems [10]. Thus, the *fluxweb* package offers the *stability.value* and *make.stability* functions using the concept of resilience to quantify the stability of a network with fluxes (see Supporting Information II for more explanations and the mathematical derivation). The second functionality provided is a sensitivity analysis of outputs regarding the parameters. The *sensitivity* function allows one to assess how the outputs of functions from the package are sensitive to a specified parameter.

## 5 Using *fluxweb*

The package can be installed from CRAN using the *install.packages*(’*fluxweb*’) command and more information is accessible on CRAN at https://bit.ly/20C0uKF. Devlopment version is available on Github at https://bit.ly/2Dkl417. Within the *fluxweb* package, we provide three complete case studies corresponding to different levels of trophic complexity (fig 1). The first example consists of a network of 62 nodes resolved to the species level and 573 edges depicting trophic interactions among soil mesofauna in a German beech forest (for details see [27]). As is often the case for species-level resolved networks, we only have a binary description of interactions (neither weight of trophic links nor feeding preferences are available). The network corresponding to the intermediate level of complexity is a version of the species-level network where species were aggregated in trophic groups using a group detection method [28]. Reducing complexity by forming aggregated groups can be used to gain basic estimates of predator foraging preferences. Here preferences were estimated by the aggregation process: the foraging preference of a trophic group *j* on a trophic group *i* is defined as the number of predation links from species of group *j* on species of group *i*. The simple case corresponds to a mesocosm of four species (one resource, two herbivores and a consumer of the two herbivores) assembled from the Chesapeake Bay river estuary [24]. Data used for the species-level food web, the group-level food web and the simple case can be accessed using the *species.level, groups.level* and *simple.case* lists respectively. Each of these lists contains all the necessary information to estimate fluxes. They are automatically loaded with the package.

**Figure 1:**
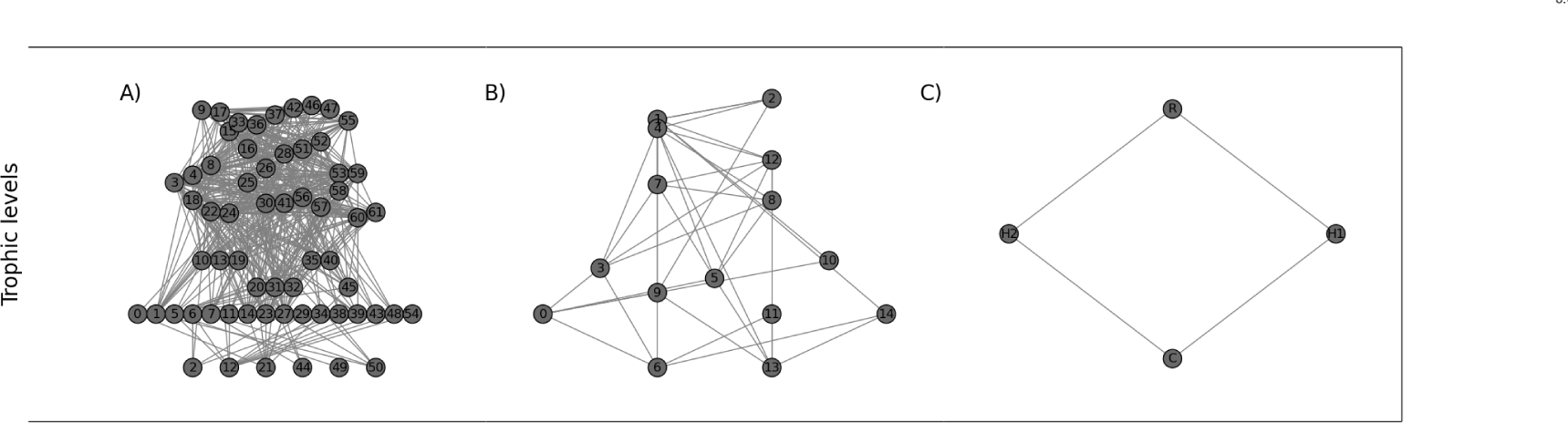
Representation of the *species.level* (A), *groups.level* (B) and *simple.case* (C) food webs.

### Species-level network

The different parameters of this dataset are:

- The network binary adjacency matrix: value of line *i* and column *j* is 1 if species *j* feeds on species *i*, 0 otherwise: `species.level$mat`
- The vector of total species biomasses (g): `species.level$biomasses`
- The vector of average species body masses (g): `species.level$bodymasses`
- The vector of assimilation efficiencies: `species.level$efficiencies`

We used species metabolic rates to define energetic losses related to physiology (eq. 8), with parameter *x*_0_ = 0.71 and *b* = –0.25 according to [4]. Values obtained are in joules per second and then scaled to joules per year.

### Group-level network

Data provided here are similar to the ones used for the `species.level` list. Body mass of a group is defined as the average bodymass of species belonging to this group. group biomass is defined as the sum of species biomass from the group. In addition, the list contains the `species.tgs` data frame indicating the identity of species in each trophic group.

### Simple-case network

In this specific case, metabolic rates are not estimated based on regressions with body masses but, as is often possible from micro‐ or mesocosm experiments, directly measured by 0_2_ respiration (*ml.mg*^-1^.*h*^-1^) and converted to joules per year. Thus, this dataset departs from the two others as no information about body mass is needed to estimate fluxes. The *simple.case* list contains:

- The network binary adjacency matrix: value of line *i* and column *j* is 1 if species *j* feeds on species *i*, 0 otherwise: `simple.case$mat`
- The vector of species biomasses (g): `simple.case$biomasses`
- The vector of species metabolic rates (j.year): `simple.case$metabolic.rates`
- The vector of assimilation efficiencies: `simple.case$efficiencies`

### *Fluxweb* function examples

The different datasets can be loaded using the `load()` function and elements can thereafter be directly accessed after a use of the `attach()` function. For the species and group level examples, species metabolic losses (per unit of biomass) have to first be estimated with eq. 8:

~~~
metabolic.rates = 0.71*bodymasseŝ-0.25
~~~

For these three cases, the matrix of fluxes is simply computed through the call to the *fluxing* function:

~~~
fluxes = fluxing(mat, biomasses, metabolic.rates, efficiencies, bioms.prefs = TRUE, bioms.losses = TRUE, ef.level = ‘prey’)
~~~

Here, `bioms.prefs = TRUE` specifies that species preferences depend on prey abundances (eq. 9). The *bioms.losses* argument is set to *TRUE* to compute metabolic losses for species populations (as they are provided per unit biomass). For the example from the mesocosm experiment, as metabolic rates were directly measured, this has to be switched to *FALSE*. The *ef.level* argument is set to *prey* as efficiencies provided in these datasets depends on prey identities.

In the same way, the stability of the food web of fluxes is returned by the *stability.value* function:

~~~
stability.value(val.mat, biomasses, losses, efficiencies, growth.rates, bioms.prefs = TRUE, bioms.losses = TRUE, ef.level = ‘prey’)
~~~

with the addition of a vector of growth rates for basal species (parameter `growth.rates`), determined using the classic allometric equation (eq. 8).

## 6 From data sampling to functions

As a very simple example how to convert community data into quantitative fluxes, we propose guidelines for experimental ecologists who want to use *fluxweb* under the assumptions of the metabolic theory of ecology.

### 6.1 Preparing the data

Using *fluxweb* will require the following data:

- a matrix defining the set of **trophic interactions** between each species pair of the ecological system considered (hereafter called *food.web*).
- A vector with the average **body masses** of species (in g, hereafter called *bodymasses*).
- A vector with the total **biomass** of each population (in g, hereafter called *biomasses*).
- A vector with the **organism type** (i.e. plant, animal or detritus) of each species (hereafter called *org.types*).

Then, a vector of **metabolic types** (as defined in Table 3) of each species (thereafter called *met.types*) is not mandatory to calculate fluxes but is a set of easily accessible information that can increase the precision of metabolic rate estimations.

**Table 2:** Description of the different functions provided by *fluxweb* and their arguments. More details can be found in the help of the package.

| Function | Description | Arguments |
| --- | --- | --- |
| *fluxing* | Compute energy fluxes in networks | - Interaction matrix (including preferences if provided)<br>- Physiological losses<br>- Feeding efficiencies<br>- Species biomasses (optional) |
| *stability.value* | Return stability of the network of flux (resilience) | - Interaction matrix of fluxes<br>- Species biomasses<br>- Physiological losses<br>- Feeding efficiencies<br>- Growth rate |
| *make.stability* | Return the smallest multiplicative scalar of losses insuring network stability (i.e. producing negative resilience) | - Interaction matrix of fluxes<br>- Species biomasses<br>- Physiological losses<br>- Feeding efficiencies<br>- Growth rate |
| *sensitivity* | Compute the sensitivity of a function to an argument | - Function to analyze<br>- Parameter to analyze<br>- Interval of uncertainty for the parameter<br>- Number of replicates to use<br>- Set of parameters needed by the function |

**Table 3:** Parameter values used for the calculation of species metabolic rates depending on their metabolic types. Values from [4]

| Metabolic type | intercept($x_0$) | exponent ($b$) |
| --- | --- | --- |
| *Ectotherm vertebrates* | 18.18 | -0.29 |
| *Endotherm vertebrates* | 19.5 | -0.29 |
| *Invertebrates* | 17.17 | -0.29 |

All of these details will allow the definition the mandatory arguments needed to calculate the fluxes: the food web (the matrix *food.web*), the physiological losses (vector *losses*), and the efficiencies (vector *efficiencies*).

#### food.web

Information about the **food web** is the first parameter required by the fluxing function. It should be a matrix (thereafter called `mat`) of *n* rows and *n* columns, where *n* is the total number of species involved in the study. The order of species should be identical between rows and columns. A non-zero value at the intersection of line *i* and column *j* in the food web matrix means that predator *j* consumes prey *i*. The values used to fill this matrix can be either binary (0/1) assuming that predators’ foraging preferences on their prey are unknown, or real values, defining these foraging preferences.

#### losses

The losses parameter will be defined in this context as metabolic rates. They are calculated using the species body masses. This calculation can be achieved using eq. 8. It is possible to define the parameters of this equation depending on species **metabolic types** (see table 3 or [25]), or to use an average value. In the case of an average value, the per unit of biomass (i.e. g) metabolic rate *X* is:

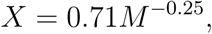

where *M* is the body mass of the species. Then, the corresponding R line of code is:

losses = 0.71 * bodymasses ̂(–0.25)

It is possible to obtain a more precise estimation of species metabolic rates, considering the parameters of Table 3 defined for each entry of the vector *met.types*. Then, the definition of the vector *losses* containing species’ metabolic rates can be achieved with:

\# first, create an empty vector where length is equal to

\# the number of species in the food web (nb.species)

losses = rep(NA, nb.species)

\# then define values associated to the different metabolic types,

\# using species ’ body mass (stored in the vector body.masses)

ecto.vert = met.types == ’Ectotherm vertebrates’

endo.vert = met.types == ’Endotherm vertebrates’

inv = met.types == ’Invertebrates’

losses [ ecto. vert ] = 18.18 * bodymasses [ ecto . vert ] ̂(– 0.29)

losses [ endo. vert ] = 19.5 * bodymasses [ endo. vert ] ̂(–0.29)

losses [inv] = 18.18 * body.masses [ inv ] ̂(—0.29)

It is important to note that the calculation of metabolic rates using the equations from the metabolic theory of ecology (8) leads to values were units are per-gram of biomass, they do not correspond to the total energetic losses of the entire populations (which can be obtained by multiplied the per-gram of biomass rates by the total biomass of the population). It is quite common in food webs to have nodes such as ’detritus’ or ’dissolved organic matter’. Values for the metabolic rates of such nodes can be set to NA if they are basal and zero in any case.

#### efficiencies

The last parameter needed to estimate fluxes is the vector of feeding **efficiencies**. Because species’ physiological losses were estimated using metabolic rates, assimilation efficiencies should be used (assimilation efficiency defines the proportion of eaten biomass that can be used for biomass production plus metabolism [26]). These efficiencies can be defined using basic information on organism types. Indeed, the efficiency with which a predator will assimilate energy from a prey can be defined by the type of prey eaten. Considering a vector *org.type* defining the organism types of food web nodes as ’animal’, ’plant’ or ’detritus’, efficiency values for these three categories are respectively 0.906, 0.545 and 0.158 [26]. The vector of *efficiencies* can be created like:

\# first, create an empty vector where length is equal to

\# the number of species in the food web (nb.species)

efficiencies = rep(NA, nb.species)

\# then define values associated to organisms types

efficiencies [ org. t ype == ’animal’] = 0.906

efficiencies [ org. t ype == ’plant’] = 0.545

efficiencies [ org.type == ’detritus’] = 0.158

### 6.2 Calculating fluxes

Once the data set is prepared as described above, the calculation of fluxes is straightforward. It is simply achieved using the fluxing function:

mat.fluxes = fluxing(mat, biomasses, losses, efficiencies)

where *mat.fluxes* is a matrix containing the fluxes between each species pair. At this point it is important to realize that we used the default behaviour of the *fluxing* function and that several options are hidden so far.

Indeed, we use the default values of the optional arguments:

- *bioms.pref = TRUE* will scale the species diet preferences (i.e. the values from the food web matrix *mat*) to the biomasses of their prey, according to this equation:

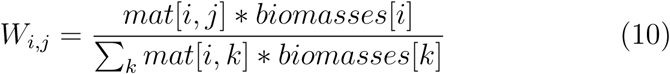

where *W*_*i,j*_ is the scaled preference of predator *j* on prey *i*
- *bioms.losses = TRUE* will calculate the total losses of species as the product of the term by term product of the vectors *losses* and *biomasses*. Thus, setting this option to TRUE corresponds to a dataset were species’ metabolic losses where defined per unit of biomass. If species losses where directly measured at the population scale (using some respiration measurement for example), this parameter should be set to FALSE.
- *ef.level = ”prey”* will assume that the species efficiencies are defined according to prey (i.e., for each species, it is the efficiency with which it will be assimilated once it has been preyed upon).

Using this methodology to compute fluxes with the *species.level* example (fig. 2) dataset would lead to the following lines of code:

\# first attach the dataset

attach (species.level)

\# create the vector of metabolic rates.

\# In this example it is done using the general allometric equation: met.rates = 0.71 * species . level $bodymasses ̂ –0.25

\# The efficiencies are already defined in the efficiencies vector.

\# Then the network of fluxes is obtained using:

mat.fluxes = fluxing(mat, biomasses, met.rates, efficiencies)

**Figure 2:**
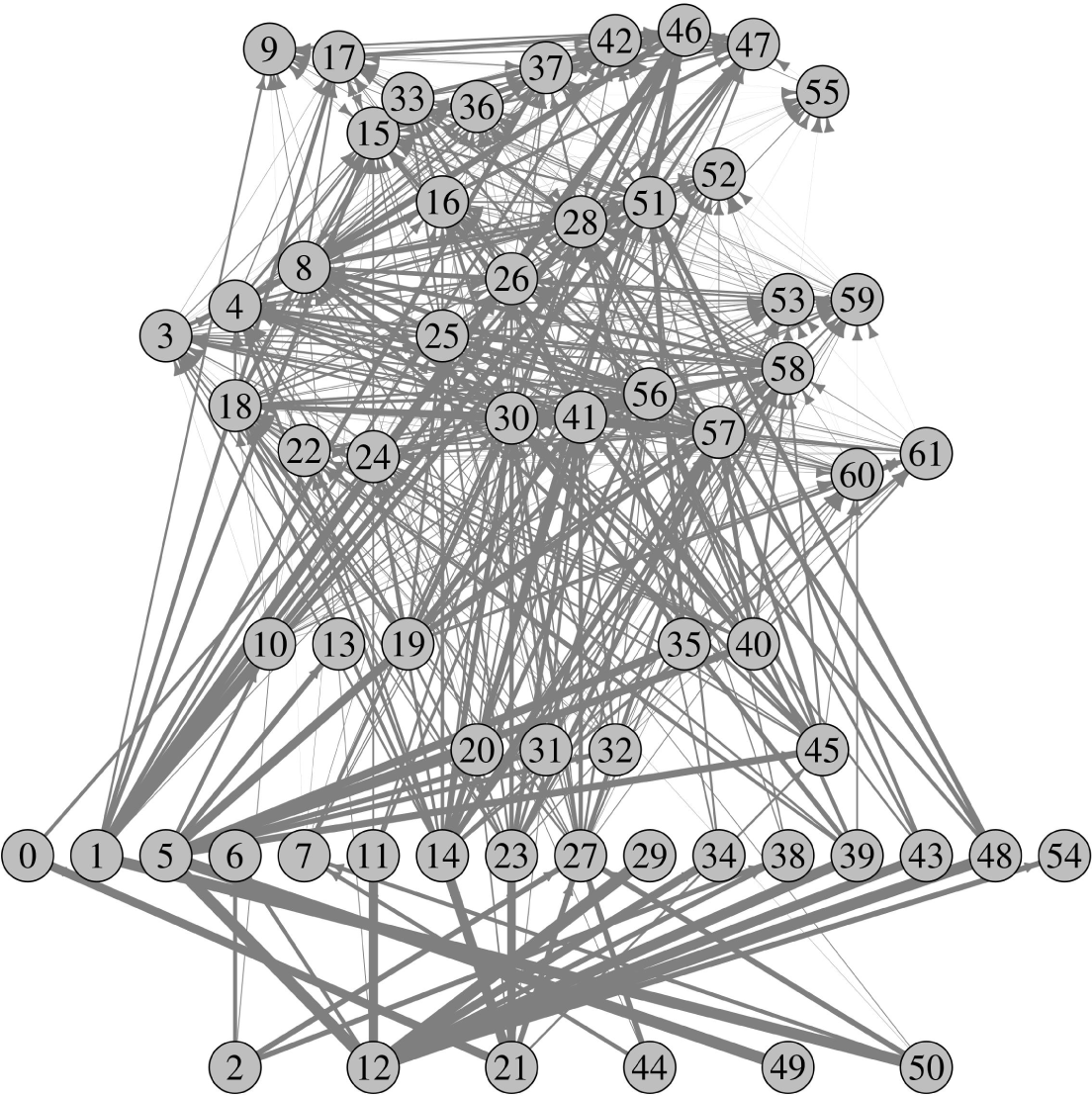
Representation of the *species.level* food web. Width of links scales with the log of fluxes. Nodes’ labels correpond to the species ordering in the *species.level* dataset

### 6.3 From fluxes to function

Once the matrix of fluxes is obtained, it is possible to estimate some ecosystem functions such as herbivory, detritivory or carnivory. In the following, we will define them as the sum of fluxes outgoing from plant, detritus and animal nodes, respectively. It is important to note that the fluxes estimated by the *fluxing* function correspond to energy loss from resource nodes. They differ from the energy assimilated by consumer nodes due to assimilation efficiencies. Thus, functions from the species.level example (fig. 3) can be estimated by simple sum operations on the *mat.fluxes*:

\# basal species are species without prey

basals = colSums (mat.fluxes) == 0

names [basals ]

\# plants are basal species that are not organic matter or exudates

plants = basals

plants [which (names == ’dead organic matter’

| names == ’root exudates’)] = FALSE

\# Herbivory is defined as the sum of fluxes

\# outgoing from plant consumers

herbivory = sum(rowSums (mat .fluxes [ plants, ]))

\# Carnivory is defined as the sum of

\# fluxes outgoing from animals

carnivory = sum(rowSums (mat .fluxes [! basals, ]))

\# detritivory is defined as the sum of fluxes

\# outgoing from detritus consumers

detritivory = sum(mat .fluxes [names == ’dead organic matter’ ,])

\# total fluxes

total = sum(mat.fluxes)

**Figure 3:**
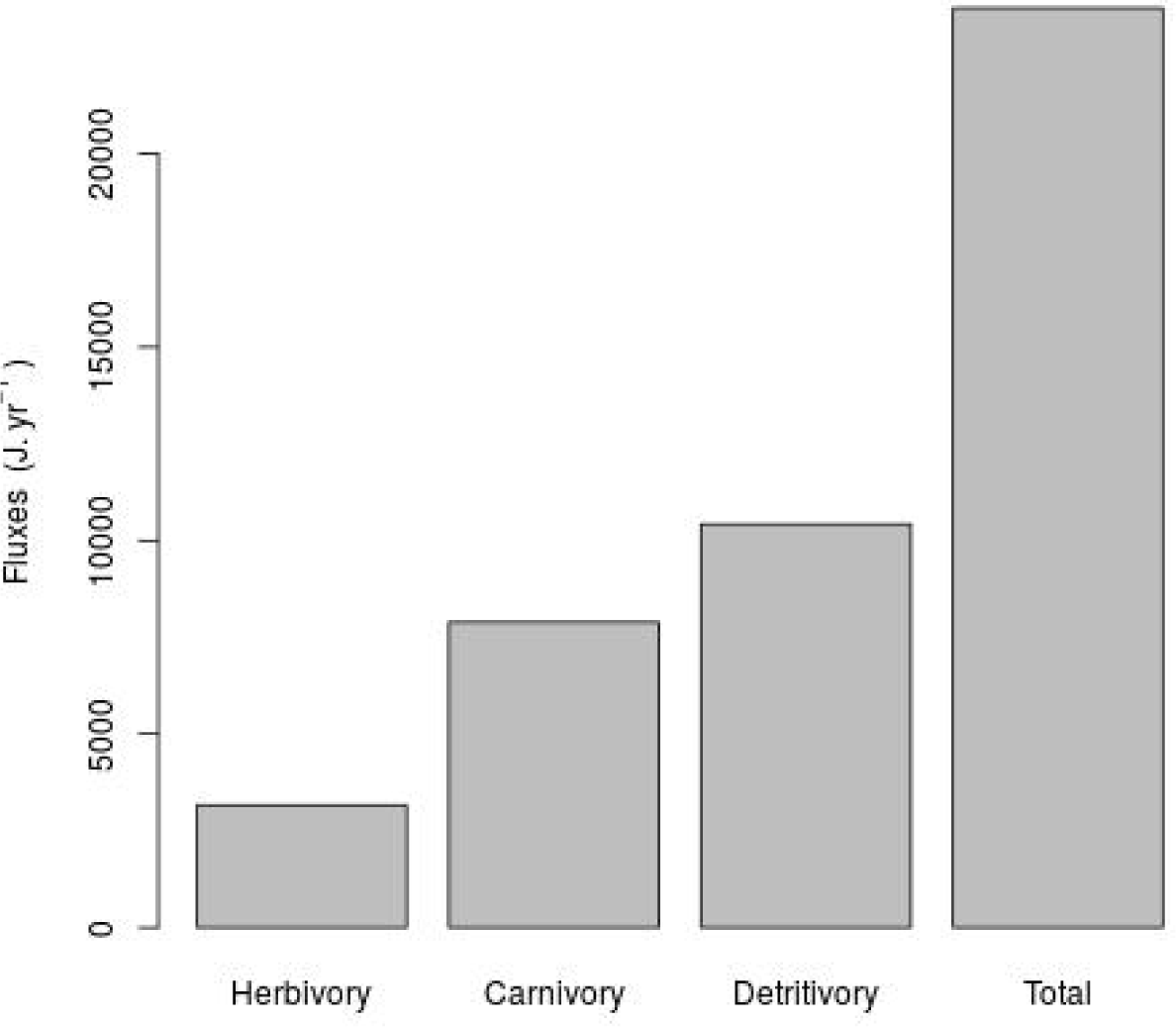
Estimation of the herbivory, carnivory and detritivory functions for the *species.level* food web, as well as the total amount of energy transiting in the food web over one year.

## 7 Conclusion

The R package *fluxweb*, provides a set of methods allowing the calculation of energy fluxes in food webs based on the conceptual framework of the ‘food web energetics’ approach [2, 9, 17, 3]. Fluxes within a system, which have typically been estimated in highly aggregated compartments, can now be quickly calculated at the species level or grouped as appropriate by users to match their objectives. This provides an advance to mechanistically understand how changes in biodiversity may impact ecosystem functioning [6], and is timely given the increasing amount and complexity of ecological network data being collected over environmental and disturbance gradients [29, 30]. Under the assumption of biomass equilibrium, multiple aspects of ecosystem function can be investigated owing to the package’s flexibility in the processes represented by parameters, their units, and how the outputs are interpreted. Function parameters can be estimated from general allometric relationships suitable for abstract models or tuned accordingly to precise measurement of specific systems depending on the users’ aims and on the availability of project-specific measurements or system-specific literature values. It is also possible to drop the hypothesis of equilibrium in case of the use of temporal dataset by adding changes of species biomass in time to the loss parameter. The impact of these estimations on ecological inferences can be assessed with the sensitivity function (Supporting Information III).

Several ecosystem functions can easily be estimated. For example, primary production can be defined as the sum of fluxes outgoing from plant species [31] (because outgoing fluxes from plants must be balanced by ingoing fluxes, thus providing an indication for total uptake by plants). Hypotheses regarding the effect of network structure or community composition on a single function (or multiple single functions; multifunctionality) can also be tested, such as secondary production by herbivores or decomposition by detritivores [9, 3]. Assessing such fluxes is important because they are directly linked to ecosystem services but may be mismatched with the standing-stock biomass of these species or trophic groups [9]. Additionally, whole-system flux, the sum of the entire fluxing matrix, can be used as a single value representing the emergent property of multitrophic functioning [9].

The functions of *fluxweb* also offer several distinct but related ways to examine network stability that are important in the face of global changes and species loss. First, the biomass fluxes can be interpreted as link weights, and used to assess the distribution of interaction strengths in the network. Second, the stability function returns the network resilience, its ability to return to its equilibrium state following a small perturbation (see Supporting Information II). Overall, the *fluxweb* package thus offers important tools for research on quantitative food webs and ecosystem functioning in real and dynamic natural landscapes [32].

## Acknowledgements

M.J., D.G. and A.B. were supported by the German Research Foundation within the framework of the Jena Experiment (FOR 1451). MJ acknowledge the Swiss National Science Foundation. U.B., A.B., D.G., J.H., B.R., S.W. and BG gratefully acknowledge the support of the German Centre for Integrative Biodiversity Research (iDiv) Halle-Jena-Leipzig funded by the German Research Foundation (FZT 118). BG thanks Stephane Legendre for helpful discussions at the beginning of this project.

## 8 Supplementary information I - mathematical resolution

### 8.1 Efficiencies depending on predator identity

We will consider in the following that feeding efficiencies depend on predator identity. We define *e* as the vector of efficiencies and *W* as the matrix such that *W*_*ij*_ is the proportion of energy entering *j* that is obtained from *i* (Σ_*j*_*W*_*ij*_ = 1). *F*_*ij*_ is the flux from species *i* to i. *L*_*i*_, the energy loss of species *j* is defined by:

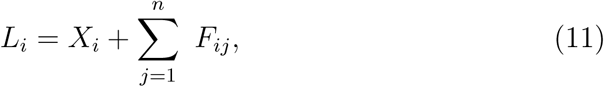

where *n* is the number of species and *X*_*i*_ are the physiological losses of species *i*. Thus, for satisfying the equilibrium criteria, *F*_*i*_, the sum of fluxes entering *i* is:

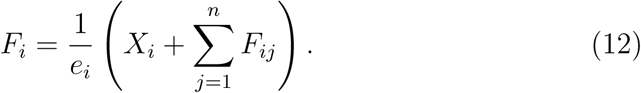

As *W*_*ij*_ sets the proportion of energy entering *j* obtained from species *i*, using *F*_*ij*_ = *W*_*ij*_ *F*_*j*_, we can write

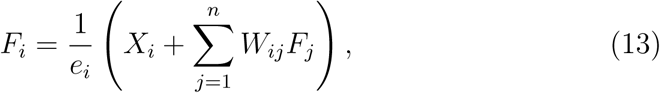

where values *W*_*ij*_ are estimated accordingly to species preferences (*w*_*ij*_) and prey abundances:

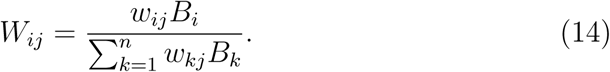

We then have:

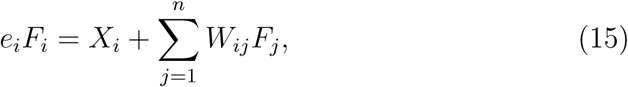

which can be rewritten as:

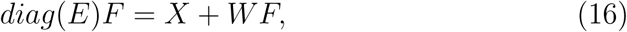

where *diag(e*) is the diagonal matrix such that *diag(e*)_*ii*_ = *e*_*i*_. Provided that (*diag(e*) – *W*) is invertible, the system solves as:

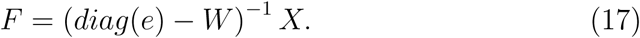

Then, all fluxes *F*_*ij*_ = *W*_*ij*_*F*_*j*_ are derived from *F*_*j*_ using *W*.

### 8.2 Efficiencies depending on prey identity

Another common method is to define feeding efficiencies according to prey identity. This section proposes a method to adapt the previous framework to this case.

As preferences are defined at the prey level, we need to adapt the previous framework by adding a nutrient node on which all basal species feed with an efficiency of one. Then, eq.4 becomes:

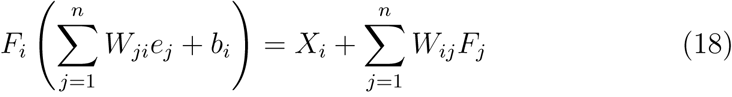

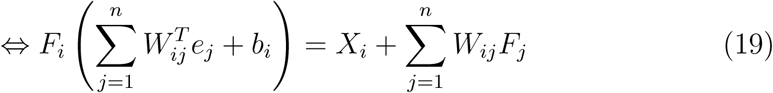

were *b*_*i*_ is 1 if *i* is a basal species, 0 otherwise. This can be rewritten as:

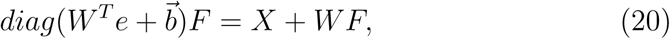

and, provided that 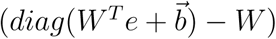 is invertible, solved by:

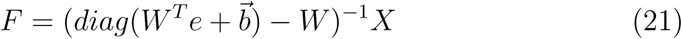

### 8.3 Efficiencies depending on the link identity

It is possible to generalise this approach to efficiencies defined for each prey-predator couple. The solution needs the definition of matrix *U* such as *U*_*ij*_ = *W*_*ij*_*e*_*ij*_. Then, eq. 4 becomes:

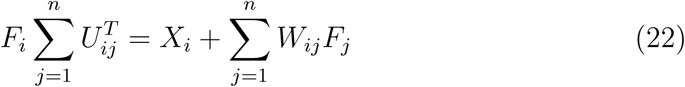

and the system then reads:

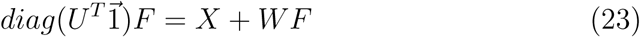

where 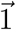 is the vector of ones. System is solved as:

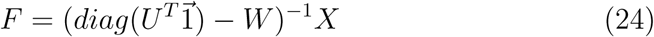

## 9 Supplementary information II

This document presents how the fluxes calculated under the steady state hypothesis can easily be used to assess system stability, following the framework of Moore and De Ruiter [21]. Here we use resilience as a definition of stability. Resilience is determined from the Jacobian matrix. The system is in a stable equilibrium only if the real parts of eigenvalues from the Jacobian are all negative. In this case, resilience is the absolute value of the real part of the largest eigenvalue, which is the value returned by the *stability* function from the *fluxweb* package.

Another measure of stability, provided by the function *make.stability*, is to find the minimal value of a scalar s defining the proportion of physiological losses related to species density. In this case, physiological loss terms in the diagonal of the Jacobian matrix are now defined as *sX*_*i*_ and directly affect the resilience value, *s* being the measure of stability. We will show in the following section how fluxes at equilibrium can relate to a Lotka-Volterra system in an equilibrium state, and how to compute the Jacobian matrix, first assuming that feeding efficiencies relate to predator identity and then assuming that they depend on prey identity.

### 9.1 Derivation of the Jacobian matrix

#### 9.1.1 Preferences defined at predator level

We can consider the following system of equations, describing the dynamics of population biomasses in a community:

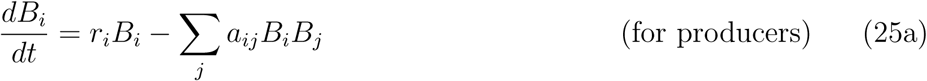

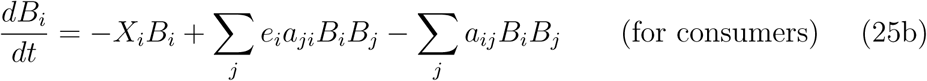

*a*_*ij*_ is the coefficient of interaction between prey *i* and predator *j* and *r*_*i*_ is the relative growth rate of producer *i*. *P*_*i*_ and *p*_*i*_ respectively define the sets of predators and prey of species *i*. This model assumes a type I functional response *f*_*ij*_ defined as:

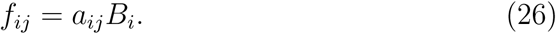

As the whole method assumes that fluxes and biomasses are at an equilibrium state, we have:

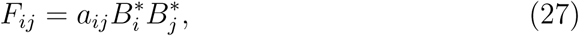

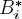 denoting biomass of species i at equilibrium. Then, off-diagonal elements *α*_*ij*_ from the Jacobian matrix correspond to the per capita effects (effect of one unit of species biomass). Considering the possible presence of cycles of length 1 (species *i* is at the same time a prey and a predator of species *j*), off diagonal elements are

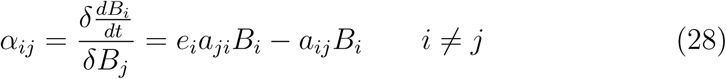

and at equilibrium, from eq. 27 we have 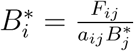 and 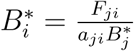. We can use it to replace elements from eq. 28 and obtain:

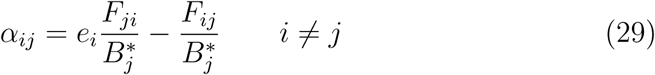

Diagonal elements, considering possible cannibalistic loops, for producers (*p*) and consumers (*c*) are:

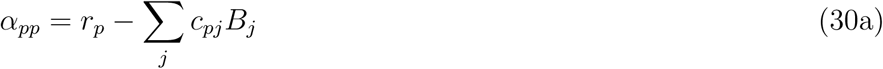

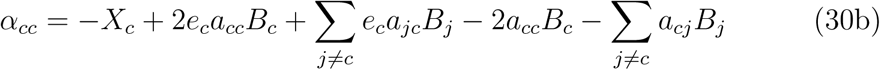

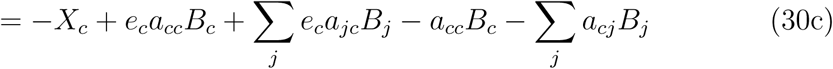

with *a*_*ii*_ ≈ 0 only if species *i* is cannibalistic. Again, using 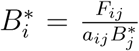 and 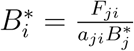 we obtain at equilibrium:

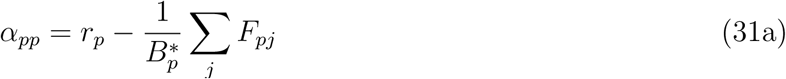

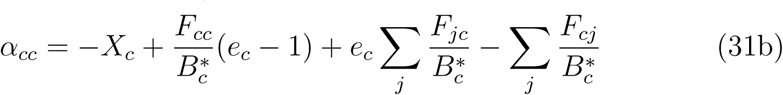

We can observe here that *α*_*cc*_ can be rewritten as

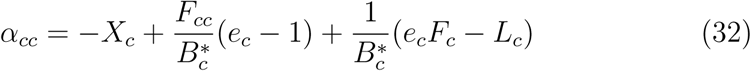

where *F*_*i*_ sets the sum of fluxing ingoing to species *i* and *L*_*i*_ its losses due to consumption. As we assume a steady state, ingoing fluxes compensate outgoing fluxes plus physiological losses: *e*_*c*_*F*_*i*_ = *L*_*i*_ + *X*_*i*_. From that, we obtain:

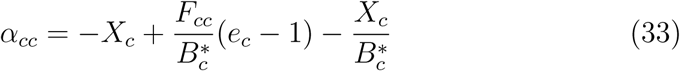

#### 9.1.2 Efficiencies defined at prey level

The Lotka Voltera system is now written as:

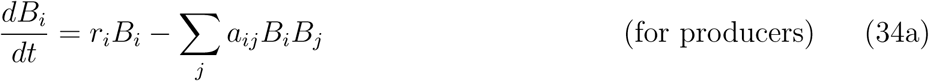

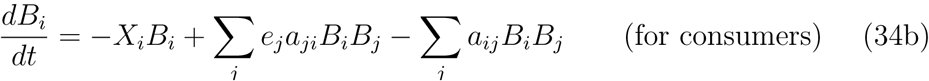

Here *e*_*j*_ defines efficiency of prey species *j*. At equilibrium, off-diagonal elements of the Jacobian are as above:

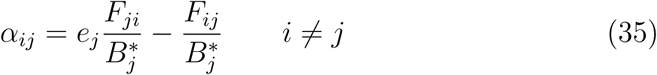

Diagonal elements, considering possible cannibalistic loops, for producers (*p*) and consumers (*c*) are:

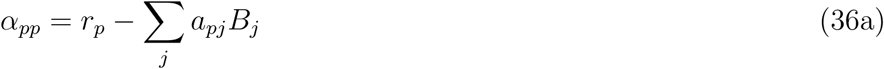

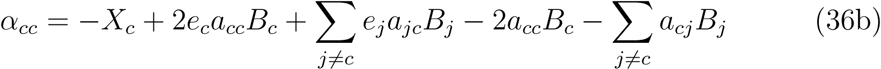

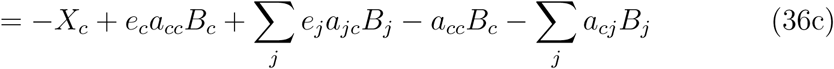

which, at equilibrium leads, like above, to:

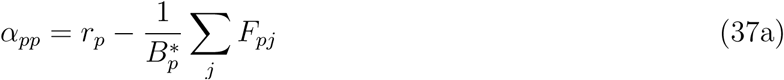

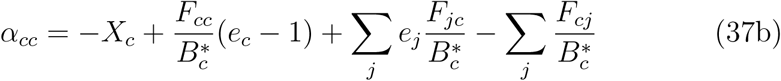

#### 9.1.3 Preferences defined at link level

Following the same mathematical derivation as before, we obtain:

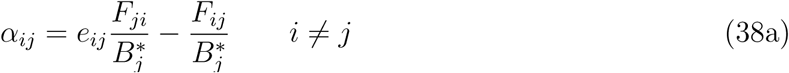

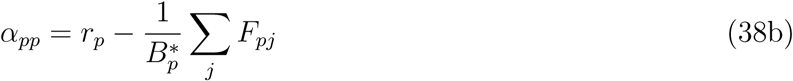

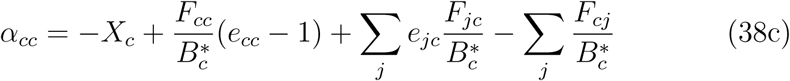

## 10 Supplementary information III - sensitivity to input parameters

We estimated here if the uncertainty or the lack of precision of the estimation of parameters tended to lead to large errors in the estimation of fluxes. To do so, we used the *sensitivity* function to estimate the sensitivity of the *fluxing* function to input parameters. The *sensitivity* function applies a random variation to a selected input parameter of the *fluxing* function. As a result, it returns a matrix containing, for each for each flux, its average coefficient of variation, estimated as:

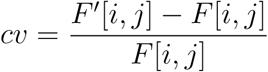

were *F*[*i*, *j*] is the flux from species *i* to species *j* when no variation is applied to parameters and *F*′[*i*, *j*] is its equivalent when a random variation is applied.

Here, we considered the sensitivity of the *fluxing* function to the *losses*, *efficiencies* and *preferences* parameters using the *species.level* example.

For each of these parameters, we simulated an uncertainty on the precision of the estimation method by increasing the value of the *var* parameter of *sensitivity* from 0 to 0.12 (by steps of 0.01) using 50 replicates each. Thus, for each flux and each parameter variation, we obtained the standard deviation of its departure (*cv*) to the original value. To summarise these results, we calculated the mean of these standard deviations over all fluxes for each parameter variation. We then obtained a scalar value representative of the uncertainty of the result of the *fluxing* function depending on the lack of precision of parameter estimation. The fig. 4 was generated using the following code:

attach (species.level) set.seed(12)

losses = 0.71 * bodymasses ′–0.25

\# creation of vectors to store the standard deviation of c.v.

\# for each uncertainty level

sd.cvs.eff = c ()

sd.cvs.los = c ()

sd.cvs.mat = c()

for (var in seq(0, 0.12, 0.01)){

cat (’ var : ’, var, ’ \n ’)

\# for efficiencies

res = sensitivity (fluxing, ” efficiencies ”, var, 50,

mat = mat ,

biomasses = biomasses ,

los ses = losses ,

efficiencies = efficiencies)

sd.cvs.eff = c(sd.cvs.eff, mean(res [[2]], na.rm = T))

\# for losses

res = sensitivity (fluxing, ’’losses”, var, 50,

mat = mat ,

biomasses = biomasses ,

loss es = losses ,

efficiencies = efficiencies)

sd.cvs.los = c(sd.cvs.los, mean(res [[2]], na.rm = T))

\# for preferences

res = sensitivity (fluxing, ”mat”, var, 50,

mat = mat ,

biomasses = biomasses ,

loss es = losses ,

efficiencies = efficiencies)

sd.cvs.mat = c(sd.cvs.mat, mean(res [[2]], na.rm = T))

}

plot (s d . c vs . eff ~ seq, xlim = c(0,0.12),

xlab = ’ variation in parameters ’ ,

ylab = ’ observed departure to original results ’ ,

pch = 16)

point s (s d . cv s . lo s ~ seq, col = ’red’, pch = 16)

points (sd.cvs.mat ~ seq, col = ’green’, pch = 16)

abline (a = 0, b= 1, lty = 2)

legend (’ topleft ’ ,

legend = c(’ efficiency ’, ’ metabolic losses ’, ”species ’ ’preferences”),

col = c(’ black ’, ’ red ’, ’ green ’) ,

pt.cex=1.5, bty=’n’,

pt.bg = c (’ black ’, ’red’, ’green’), pch = 21)

**Figure 4:**
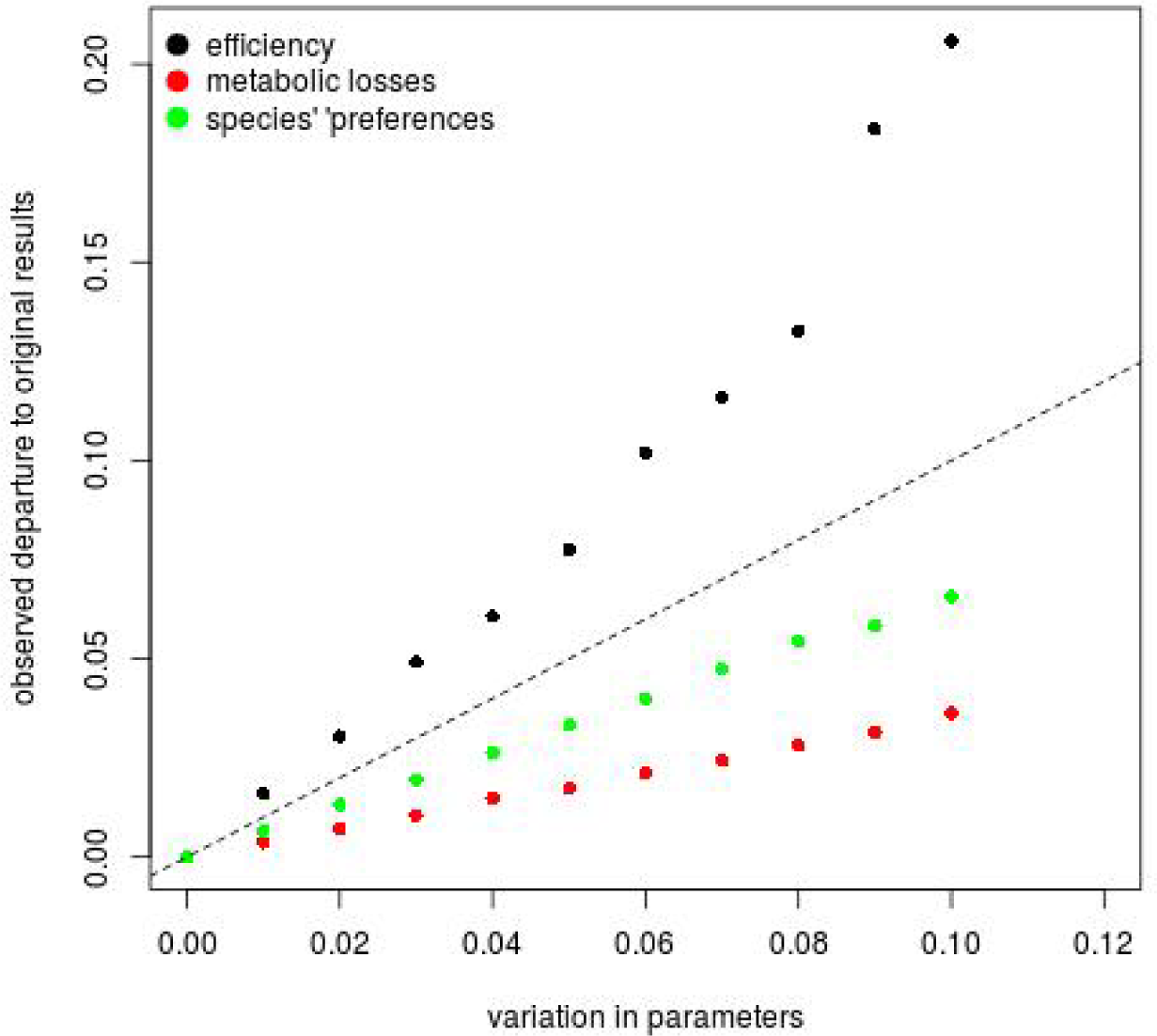
Representation of output uncertainties of the *fluxing* function (y axis) when a random variation is applied to an input parameter. The x axis represent the uncertainty applied to parameters (*var* argument of the *sensitivity* function). The dashed line represents the identity.

